# CRM1 regulates androgen receptor stability and impacts DNA repair pathways in prostate cancer, independent of the androgen receptor

**DOI:** 10.1101/2024.02.13.579966

**Authors:** Rajendra Kumar, Janet Mendonca, Abhishek Shetty, Yuhan Yang, Olutosin Owoyemi, Lillian Wilson, Kavya Boyapati, Deven Topiwala, Naiju Thomas, Huong Nguyen, Jun Luo, Channing J. Paller, Samuel Denmeade, Michael A. Carducci, Sushant K. Kachhap

## Abstract

Among the known nuclear exportins, CRM1 is the most studied prototype. Dysregulation of CRM1 occurs in many cancers, hence, understanding the role of CRM1 in cancer can help in developing synergistic therapeutics. The study investigates how CRM1 affects prostate cancer growth and survival. It examines the role of CRM1 in regulating androgen receptor (AR) and DNA repair in prostate cancer. Our findings reveal that CRM1 influences AR mRNA and protein stability, leading to a loss of AR protein upon CRM1 inhibition. Furthermore, it highlights the involvement of HSP90 alpha, a known AR chaperone, in the CRM1-dependent regulation of AR protein stability. The combination of CRM1 inhibition with an HSP90 inhibitor demonstrates potent effects on decreasing prostate cancer cell growth and survival. The study further explores the influence of CRM1 on DNA repair proteins and proposes a strategy of combining CRM1 inhibitors with DNA repair pathway inhibitors to decrease prostate cancer growth. Overall, the findings suggest that CRM1 plays a crucial role in prostate cancer growth, and a combination of inhibitors targeting CRM1 and DNA repair pathways could be a promising therapeutic strategy.

## Introduction

Extensive evidence suggests that the nuclear export machinery in many cancers is dysregulated to potentiate cancer development and progression. Dysregulated nuclear export can result in mislocalization of specific proteins; notably, such mislocalization can inactivate tumor suppressor function and activate oncogenic gene function [1, 2]. CRM1 (also called Exportin1) is the most studied prototype among the seven known exportins present in mammalian cells. CRM1 and RanGTP bind to specific nuclear export motifs present in the cargo proteins to export them out of the nucleus [3, 4]. CRM1 exports nearly 700 cargo proteins outside the nucleus, including several DNA repair proteins, which include key tumor suppressors such as p53 and BRCA1, among others [2, 5]. Our work was one of the earliest to demonstrate that CRM1 can be successfully targeted in prostate cancer using selective inhibitors of nuclear export (SINE) [6]. Our study indicated that androgen receptor-positive prostate cancer cells were highly sensitive to CRM1 inhibition compared to androgen receptor-negative prostate cancer cells. Another observation from this study was that prostate cancer cell death by CRM1 inhibition was potentiated by doxorubicin, a DNA-damaging agent. Others have found similar synergism with CRM1 inhibition and DNA damaging chemo- and radiotherapeutics [7–10]. Although CRM1 is known to regulate the nuclear export of various DNA repair enzymes and proteins, the exact role of CRM1 in major DNA repair pathways remains understudied. In prostate cancer, the androgen receptor plays an important role in the transcriptional regulation of DNA repair genes [11, 12]. However, whether inhibition of CRM1 transcriptionally impacts DNA repair via the androgen receptor is unknown. Using DNA repair and protein recruitment assays to delineate the role of CRM1 in DNA repair, we demonstrate that CRM1 may have AR-dependent and AR-independent effects on DNA repair.

## Methods

### Cell culture

LNCaP, LAPC4, PC3, and DU145 cells were cultured in phenol red-free-RPMI (Gibco, Thermo Fisher), and HEK293T cells were cultured in DMEM-high glucose (Sigma) supplemented with 10% FBS (Gemini Bio). VCaP cells were grown in DMEM media (ATCC) containing 1.5 g/L sodium-bi-carbonate. All cell lines were tested for mycoplasma contaminants using a PCR-based detection kit (Agilent). LN-95 AR knockout cells were provided by Dr. Jun Luo’s laboratory and were cultured in phenol red-free RPMI with 10% charcoal-stripped FBS.

### HSP90 CRISPR knockout generation

To generate the HSP90 KO LNCaP and LAPC4 cells, a tetracycline-inducible *cas9* expression vector (pCW-Cas9, addgene # 50661) was used to create stable Cas9 expressing lines. sgRNA against HSP90-α was cloned in pLXsgRNA vector (addgene# 50662). Lentiviral particles for cas9 and HSP90-alpha were produced using HEK293T cells by con-transfection of transgene plasmid and pMD2.G (Addgene# 12259) and psPAX2 (Addgene #12260). LNCaP and LAPC4 Cas9 expressing cells were infected with lentiviral particles and treated with 1 µg/mL doxycycline to induce cas9, followed by 10 µg/mL blasticidin (Gibco). After blasticidin selection, cells were transferred to 96 well plates to generate single-cell clones. Western blots verified knockouts in LNCaP and LAPC4 cells were expanded and cryopreserved for future experiments.

### Quantitative real-time PCR

A trizol-based (Thermo Fisher) RNA precipitation method was used to isolate the total RNA from cells using the manufacturer-recommended protocol. RNA was reversed transcribed to cDNA using an iSCRIPT cDNA synthesis kit (BioRad). qPCR was performed on the samples using iTaq Universal SYBR Green Supermix (Biorad) and gene-specific primers on a thermocycler equipped with measuring fluorescence (BioRad). Data was analyzed using ΔΔCt method after normalizing with housekeeper gene transcripts and vehicle-treated control. Primer sequences are provided in the supplementary table.

### Western blotting

Cell lysates were prepared using a 1% triton X-100 containing lysis buffer (CST) supplemented with protease inhibitor cocktail (Roche) on ice for 15 min. Lysates were cleared by centrifuging at 14000 rpm for 10 min at 4 °C. Lysates were denatured by adding 2X laemmli buffer (BioRad) and boiling at 95 °C. Samples were electrophoretically resolved using 4-15% SDS-polyacrylamide gel and were immunoblotted on a methanol-activated PVDF membrane. Post transfer, membranes were blocked using 5% skim milk, followed by incubation with primary antibodies overnight at 4°C and secondary antibodies at RT for 1hr. Blots were developed using either ECL Western blot detection reagent (GE Healthcare) for abundant proteins or SuperSignal West Femto (Thermo) for moderate or less abundantly present proteins. All the antibodies used are listed in the supplementary table 1.

### Immunofluorescence

For confocal microscopy, sterile coverslips were placed in the culture wells, and cells were grown in low density and treated or kept as the untreated control. After treatment duration, cells were fixed permeabilized using −20 °C chilled methanol in a freezer followed by 10% natural buffered formalin at 37 °C. After fixation, cells were blocked with 5% BSA for 2 hr, followed by incubation with primary antibody, both at 4°C. The next day, fluorescently labeled secondary antibodies (Molecular Probes, Thermo) were added after washing away unbound primary antibodies and incubated at room temperature for at least 2 hr or 4°C overnight. After staining with secondary antibodies, cells were washed gently to remove unbound secondary antibodies and were also postfixed to stabilize the bound antibodies. Cell nuclei were counterstained with DAPI, and coverslips were mounted on ProLong Gold antifade media (Thermo). Images were acquired on a Zeiss LSM 700 laser confocal microscope, and data was analyzed using the image processing package ImageJ (NIH).

### Colony formation assay

1×10^5^ cells were plated in 6 well plates overnight and were treated with defined concentrations of single agent selinexor or with experiment-specific second drug agent for 48h. After 48h, all the cells were collected, including dead cells, and replated in either 60 mm culture dishes or a 6-well plate for long-term culture without further drug treatment. The number of cells for long-term culture was adjusted to a very thin density to allow them to form separate colonies.

### Comet Assay

1×10^6^ LNCaP cells were plated in a 60 mm tissue culture dish and, after overnight incubation, treated with either vehicle or 1 µM selinexor for 1hr. After 1 hr, 20 µM Camptothecin was spiked in the culture dishes for 1hr. After 1 hr of camptothecin treatment, media was changed to remove camptothecin; however, selinexor-treated dishes were maintained on continuous exposure to ensure CRM1 inhibition. After 2 hrs of the recovery period for DNA damage, cells were trypsinized and counted to achieve a count of 10^5^ cells per ml. A 50 µL aliquot of cell suspension was mixed with 37 °C heated 1% low melt agarose. Immediately, cells mixed with agarose were layered on comet slides (Travigen) to spread approximately 5×10^3^ cells per drop. Slides were chilled in a refrigerator to solidify agarose, and cells were lysed using lysis buffer (Travigen). After overnight lysis at 4 °C, nuclei were incubated with an alkaline detangling buffer for 1 hr, followed by electrophoresis in an alkaline buffer. After electrophoresis, slides were washed with water twice, fixation in 70% ethanol and allowed to dry. Dried slides were stained with SYBR Gold before fluorescent microscopy. Images were analyzed using OpenComet plugin version 1.3.1 in ImageJ (NIH). Multiple parameters of DNA in the comet tail were derived. The tail moment was plotted on a violin plot to compare the control against selinexor-treated cells.

### Phosphotag gel analysis

Prostate tumor cells were transfected with wild-type hCRM1-Flg. After 48 hr, cells were treated with 1µM selinexor or vehicle for 1 hr followed by irradiation with 4 Gy radiation. Cells were rested for another 2 hr before collecting the whole cell lysate. Proteins were resolved on homemade SDS gel containing 5 mM Phos-Tag Polyacrylamide and 10 mM MnCl_2_ in the resolving gel. After resolving proteins on SDS PAGE gel, gels were washed with 1mM EDTA to remove excess MnCl_2._ Proteins were immunoblotted on the PVDF membrane, and membranes were probed with rabbit anti-Flg primary antibody and HRP conjugated anti-rabbit secondary antibody.

### CRM1 Co-IP for AR mRNA transcript analysis

LNCaP cells were grown on 100mm dishes and lysed using cell lysis buffer (Cell Signalling) upon 70% confluency. Immunoprecipitation of CRM1 was done using the CRM1 antibody (SCB sc-5595, 1µg) and rabbit IgG control (CST 2729s, 1µg). The lysates were incubated with the antibody at 4°C for 3 hours. Subsequently, Protein A Dynabeads (Invitrogen 10001D) were added and incubated for another 2 hours. The beads were washed one time with lysis buffer and two times with PBS. Eluted RNA was subsequently extracted from the beads by the Trizol reagent (Invitrogen). The isolated RNA was quantified, and 100ng was used for cDNA synthesis, followed by qPCR for AR using SYBR green chemistry. LNCaP cellular RNA was used as a positive control.

### Homologous recombination repair (HR) efficiency measurement analysis

1×10^6^ LNCaP cells were plated in 6 well plates overnight and were treated with different concentrations of selinexor (100-400 nM) for 2 h before transfection with an HR sensor plasmid pJustin [13]. Cells were transfected with 2 µg DNA with 5 µL Lipofectamine 2000 (Thermo). The next day, cells were trypsinized and immediately acquired on FACSCelseta (BD Biosciences). Data files were analyzed to count EGFP fluorescence-positive cells using FlowJo. Cells were also visualized on a fluorescent microscope (Nikon) to acquire the images.

### Statistical analysis

Suitable central tendency values were calculated for all the quantitatively measurable variables and used for statistical significance analysis. Normality tests were performed to check the normal distribution of the data. A parametric test was performed to compare the mean, and nonparametric analyses were used if data were found to follow the non-gaussian distribution. These analyses were performed using Prism version 9.5.1 (GraphPad Software), and a *p* ≤ *0.05* was considered significant.

## Results

### CRM1 inhibition decreases AR protein expression

CRM1 forms a ternary complex with RanGTP and shuttles proteins across the nuclear envelope [14]. To explore whether nucleocytoplasmic transport might impact the survival of patients with prostate cancer, we queried the TCGA public database for tumors that upregulate CRM1/RanGTP. Our data indicate that upregulation (1.8 fold increase in expression in tumor as compared to normal prostate) of CRM1/RanGTP mRNA significantly correlated with decreased patient survival in prostate cancer (Log Rank Test p-value=7.863e^-3^, n=240 samples) **(Fig. 1A)**. Transcriptional upregulation (> 1.5 fold) of CRM1 mRNA was found in 8-14% of prostate tumors including metastatic prostate cancer **(Fig.1B)**. Previous reports using the CRM1 inhibitor leptomycin B (LMB) indicated that CRM1 is not essential for nuclear export of the AR protein [15]. We confirmed this result in prostate cancer cells using CRM1 inhibitor selinexor **(Fig. 1C)**. Intriguingly, AR immunofluorescence signals were decreased after selinexor treatment, indicating reduced levels of AR **(Fig. 1D)**. To find whether CRM1 affects AR protein levels, we treated LNCaP and VCaP cells with increasing doses of selinexor. Treatment of the cells with selinexor led to a dose-dependent decrease in AR protein in LNCaP cells and both full-length AR and AR-V7 splice variant proteins in VCaP cells **(Fig. 1E)**. To detect whether the reduction in AR protein levels is unrelated to a decline in AR nuclear import [16], we conducted nuclear import assays in digitonin-permeabilized LNCaP cells, employing bacterially synthesized recombinant wild-type AR-GFP (gift from Dr. Paschal Bryce) [17]. In our system, CRM1 inhibition did not affect AR nuclear import **(Fig. 1F)**.

**Fig. 1:**
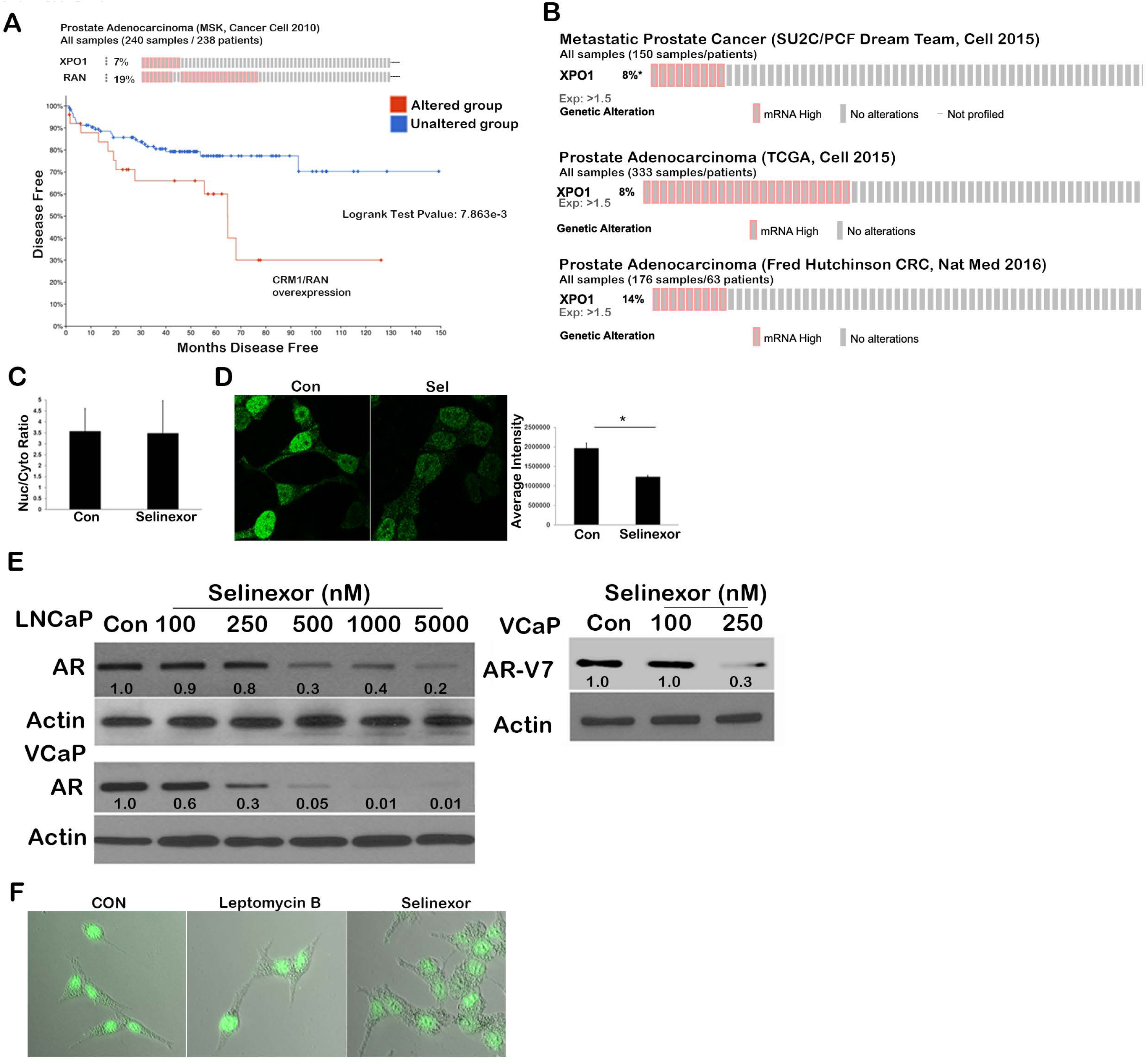
CRM1 levels in prostate cancer affect patients’ survival and alter AR levels in the PCa cell model. **A)** Survival analysis using RNA levels in patients with high or unaltered, combined CRM1 and RAN levels in primary prostate tumors (Source MSK Cancer Cell 2010, accessed through cBioPortal). **B)** Oncoprint analysis of the prostate tumor using data from 3 different data sets showing the percentage of cases with mRNA upregulation type alteration, accessed through cBioPortal. Number of patients in each dataset is mentioned on the figure. An expression cut-off value of 1.5 was used. **C)** Nuclear and cytoplasmic ratio of androgen receptor in LNCaP treated with vehicle or selinexor (1µM for 12 hours). **D)** Confocal microscopy photomicrograph showing androgen receptor levels in LNCaP cells treated with the vehicle or selinexor (dose and duration), and fluorescence intensity was calculated using ImageJ. **E)** Western blot analysis of AR (Full length (110kDa) in LNCaP and VCaP and splice variant AR-v7 (80kDa) only in VCaP cells) in selinexor-treated LNCaP and VCaP cells for 12 hrs. Actin was used for normalization, and numbers on the blot indicate the actin-normalized levels of the AR and AR-v7. **F)** Representative photomicrograph for AR nuclear import assay using recombinantly expressed GFP-fused AR. An asterisk indicates p<0.05.

### CRM1 affects AR mRNA and protein stability

In addition to protein export from the nucleus, CRM1 is known to export specific RNA subsets, including a handful of mRNAs [4, 18]. Therefore, we next asked whether AR mRNA is a CRM1 cargo RNA. We performed RNA immunoprecipitation using CRM1 antibody in LNCaP cells. Total RNA was extracted from the immunoprecipitated protein and subjected to quantitative real-time RT PCR (qRT-PCR) analysis for AR. Although we found a consistent (1 cycle) difference in AR transcripts between immunoprecipitates using CRM1 or matched IgG controls, we did not find a significant association of CRM1 with AR mRNA **(Fig. 2A)**. We next asked whether there are any proteins involved with AR mRNA transport that might serve as a CRM1 client protein and affect AR mRNA stability. The human antigen R protein (HuR) is known to help in AR mRNA nuclear export and stabilize AR mRNA [19]. HuR binds AU-rich elements in 3’ untranslated regions, stabilizing mRNA as they are exported into the cytoplasm [20]. Treatment of LNCaP cells with selinexor inhibited HuR nuclear export and restricted HuR to the nucleus **(Fig.2B)**. Restriction of HuR nuclear shuttling might affect AR mRNA stability. To investigate whether this is indeed true, we used actinomycin D, an inhibitor of *de novo* transcription, and measured AR in LNCaP cells treated with either actinomycin D or a combination of actinomycin D and selinexor at various intervals after treatment. As shown in **Fig. 2C**, compared to actinomycin-treated controls, CRM1 inhibition significantly decreased the stability of AR mRNA. To determine whether CRM1 could impact AR protein stability independently of its effect through AR mRNA stability, we treated LNCaP cells with cycloheximide, a protein translation inhibitor, in conjunction with selinexor treatment. As shown in **Fig. 2D**, selinexor treatment significantly decreased protein stability. More importantly, decreased AR levels led to a decrease in AR-mediated transcription after CRM1 inhibition, evaluated by quantitating KLK3 and Nkx3.1 expression **(Fig. 2E)**. These results suggest that CRM1 could affect both AR mRNA and protein stability and attenuate AR signaling.

**Fig 2:**
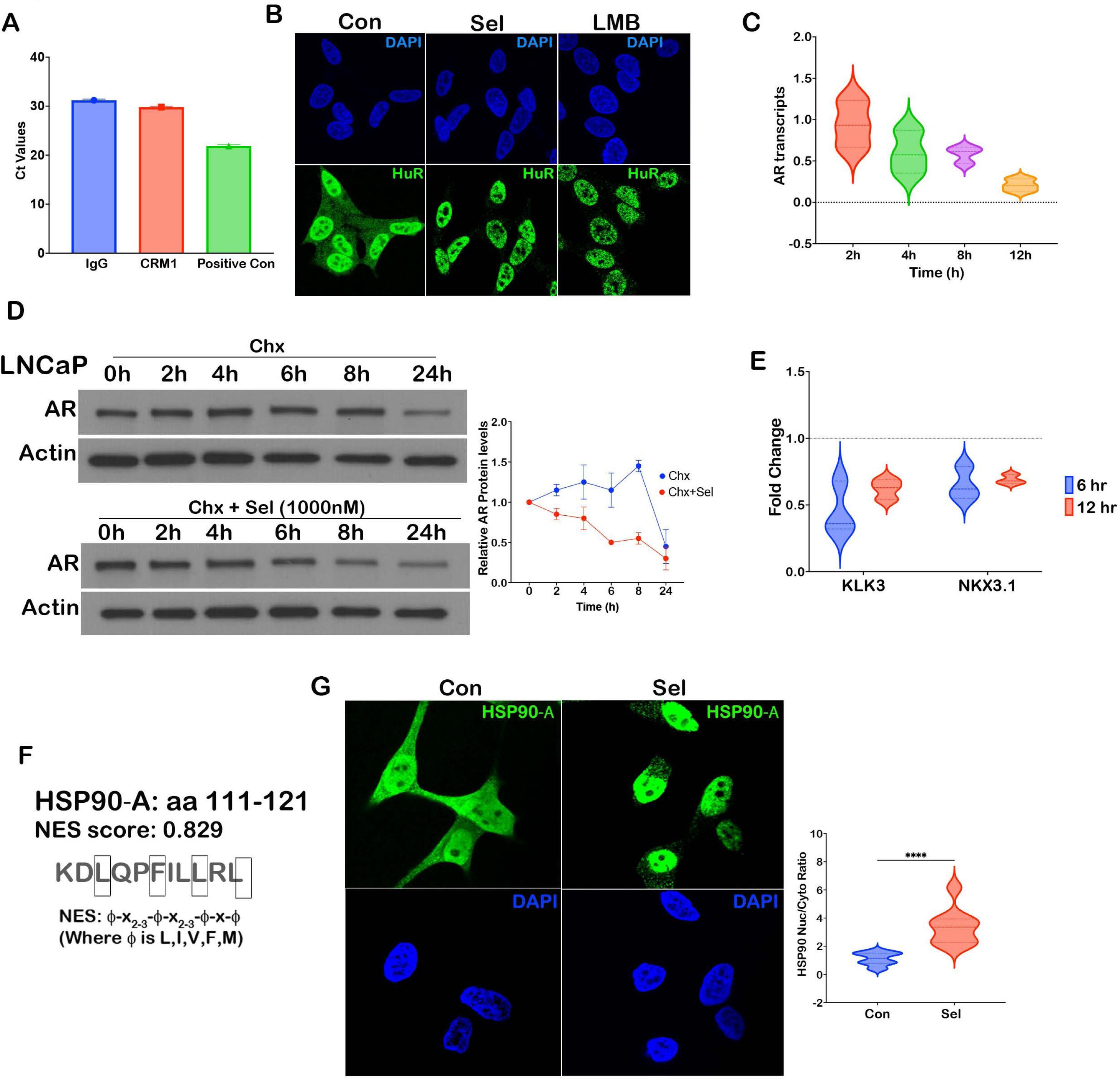
Effect of CRM1 inhibition on AR mRNA and protein stability. **A)** qRT-PCR Ct values for AR transcript from pulldown assay using control IgG or CRM1 recognizing antibody. LNCaP cDNA was used as a positive control. **B)** Localization of human antigen R protein (HuR) in the vehicle, selinexor (1µM), or leptomycin B (10nM) treated LNCaP cells for 12 hrs. **C)** Violin plot showing quantitation of AR transcripts in LNCaP cells treated with 5 µM actinomycin alone or in combination with selinexor (dose) followed by RNA isolation and qPCR analysis post-indicated time duration. **D)** LNCaP cells treated with cycloheximide (100µM) alone or in combination with selinexor (1µM) and protein samples were collected and probed with AR antibody (110 kDa). Actin (42 kDa) was used as a housekeeper protein for normalization. Actin normalized and 0hr normalized levels or AR are plotted on a line graph. The blot shows data from a representative exp of 3 independent repeats. **E)** KLK3 and NKX3.1 transcripts were measured using qRTPCR on total RNA isolated from LNCaP cells treated with 1µM selinexor for 6 or 12 hrs. **F)** HSP90α showing nuclear localization signal score. **G)** Immunostaining of HSP90α in LNCaP cells treated with vehicle or selinexor (1µM) and nuclear to cytoplasmic ratio was calculated and depicted on violin plot.

### CRM1 affects the nuclear shuttling of HSP90 alpha

How might CRM1 inhibition destabilize AR protein? We considered that CRM1 might be necessary for the function of AR-stabilizing chaperone proteins. Chaperone heat shock proteins form a multimeric complex called the foldosome, consisting of Hsp70, Hsp40, Hop, Hsp90, and p23. This complex interacts with the AR ligand binding domain (lacking in AR-V7 protein) and stabilizes the protein in a high-affinity ligand binding state [21]. Lack of these chaperone proteins can lead to misfolding and subsequent degradation of full-length AR protein [22, 23]. Some of the smaller subunits of the foldosome complex (p23, Hsp40) can freely diffuse across the nuclear pore complex without the aid of CRM1. Hence, we explored whether the cellular distribution of larger proteins in the complex, like HSP70 and HSP90, utilize CRM1 for their nuclear export. Previous studies in yeast have shown that HSP70 harbors an nuclear export signal (NES) and uses CRM1 for its nuclear export [24]. A search for NESs in the HSP90 alpha subunit indicates that it harbors a strong NES (LocNES score=0.829, **Fig. 2F**), suggesting a potential for interaction with CRM1. In congruence with this data, we found that inhibition of CRM1 in LNCaP cells by selinexor significantly increased nuclear accumulation of HSP90 protein, indicating disruption of nucleocytoplasmic shuttling of HSP90 (**Fig. 2G**).

### HSP90 inhibition potentiates CRM1 inhibition to decrease prostate cancer survival

Thus, inhibition of CRM1 may restrict HSP90 to the nucleus, precluding it from being incorporated into the cytoplasmic foldosome complex necessary to stabilize the newly formed AR. Based on this premise, we investigated whether inhibiting HSP90 will further augment the decrease in AR by selinexor. As seen in **(Fig 3A**), a combination of selinexor and geldanamycin, a potent HSP90 inhibitor, greatly decreased AR protein stability in all the AR-positive lines we tested. Encouraged by these results, we tested whether this observation would translate into cell growth inhibition. In the past, we have shown that CRM1 inhibition results in extensive apoptosis in prostate cancer cells [6]. We investigated whether treatment with a combination of geldanamycin and selinexor would result in more significant apoptosis. As seen in **Fig. 3B**, the combination greatly enhanced PARP cleavage, indicating apoptosis in AR-positive prostate cancer lines. We next investigated whether the combinatorial treatment would decrease the survival of prostate cancer lines. We treated AR-positive LNCaP and LAPC4 cells and AR-negative PC3 cells with low concentrations. From past studies [6], we know that these concentrations of selinexor did not affect PC3 cell growth, which is highly resistant to CRM1 inhibition. Clonogenic survival assays indicated that combining selinexor with geldanamycin was highly potent in decreasing clonogenic survival when low doses of geldanamycin were used **(Fig, 3C and quantified in 3D)**. This effect was not observed at elevated geldanamycin doses because, at higher concentrations, it significantly decreased survival on its own, and the combined effect could not be discerned. Surprisingly, the AR-negative PC3 cells, which are highly resistant to selinexor, also showed a greatly enhanced growth inhibition when combined with geldanamycin. These results suggest that inhibition of CRM1 in combination with HSP90 inhibition could be highly potent in decreasing the survival of both AR-positive and AR-negative prostate cancer cells. Since our results indicated that CRM1 could regulate HSP90 alpha nuclear export and since HSP90 alpha is reported to be part of the foldosome complex regulating AR stability, we investigated whether deletion of HSP90 alpha would abrogate or mute the effect of selinexor on AR protein stability. To test this, we created HSP90 alpha gene knockout in AR-positive LNCaP cells using CRISPR-Cas9 gene editing system **(Fig. 4A)**. As seen in **Fig. 4B**, treatment with cycloheximide indicated the half-life of AR protein is lower in HSP90 alpha KO (between 4-6 h) than wild type controls (between 8-24 h, **Fig. 2D**). This observation is in line with other reports suggesting that the foldosomes subunits, regulate AR protein stability. Treatment with selinexor and cycloheximide did not further enhance AR degradation but had a protective effect on AR degradation **(Fig. 4B)**. Intrigued by this observation, we first measured apoptosis in HSP90 alpha KO cells after treatment with selinexor and found that these cells were highly resistant to apoptosis even at 5µM concentration **(Fig. 4C)**. To test whether this phenomenon is not restricted to LNCaP cells, we created HSP90 alpha KO in LAPC4 cells **(Fig, 4D)**. Treatment with selinexor mirrored the apoptosis-preventive effect of HSP90 alpha KO seen in LNCaP cells. To investigate whether this translated into differences in cell survival upon selinexor treatment, we performed clonogenic survival assays on HSP90 KOs in LNCaP and LAPC4 with varying concentrations of selinexor. As seen in **Fig. 4E**, HSP90 alpha KO increased the clonogenic survival of both LNCaP and LAPC4 cells. We hypothesized that the effect could be due to increased CRM1 expression in HSP90 alpha KO cells. We evaluated CRM1 protein expression in HSP90 KO cells, and we found no change in the protein levels in KO cells as compared to wildtype controls **(Fig. 4F)**. In a previous study, we reported that treatment with selinexor results in proteasomal degradation of CRM1 [6]. We investigated whether this feature is preserved in HSP90 alpha KO cells after treating cells with increasing concentration of selinexor and probing for CRM1 protein using a western blot. We did find a decrease in CRM1 in both wild-type and HSP90 alpha KO cells in a dose-dependent manner **(Fig. 4D)**. These results suggest that while HSP90 targeting with geldanamycin (which targets both HSP90 beta and alpha) may potentiate the ability of selinexor to trigger apoptosis, cells with decreased HSP90 alpha may have adaptive signaling mechanisms that render them resistance to CRM1 inhibition.

**Fig 3:**
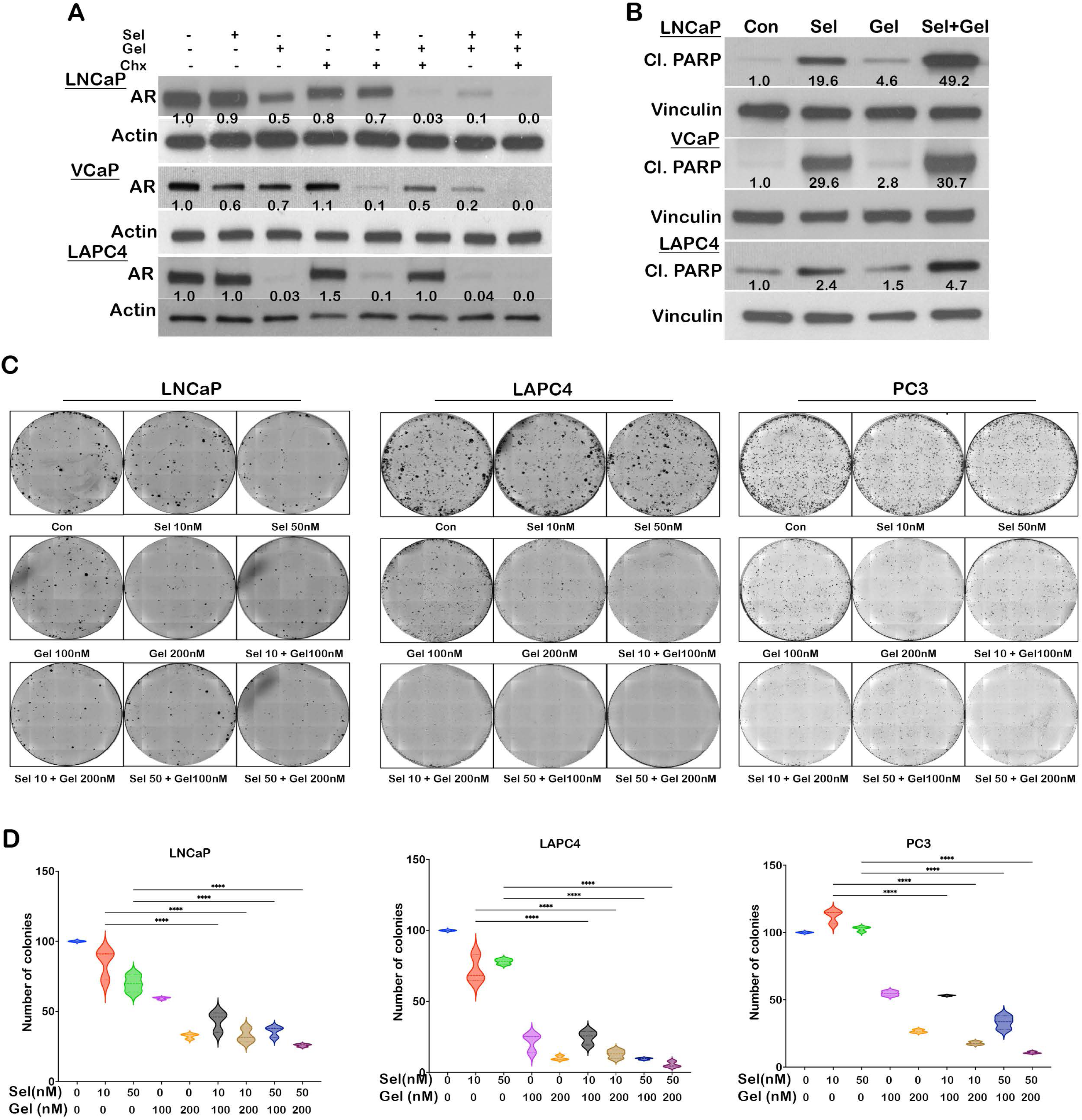
HSP90 inhibition potentiates CRM1 inhibition to decrease prostate cancer survival. **A)** Western blot showing levels of AR protein (110 kDa) in LNCaP, VCaP, and LAPC4 cells treated with vehicle, Selinexor (500 nM for LNCaP and LAPC4 and 250 nM for VCaP), Geldanamycin (400 nM), Cycloheximide (100µM) alone or in combination. Actin was used as a housekeeper protein (42kDa). **B)** Immunoblotting of cleaved PARP (89 kDa) in LNCaP, VCaP, and LAPC4 treated with vehicle, selinexor (500 nM for LNCaP/LAPC4 and 250 nM for VCaP), geldanamycin (400 nM), or combination. Actin (42 kDa) was used as housekeeper control. **C)** Representative image of colonies stained for LNCaP, LAPC4, or PC3 cells treated with selinexor (10 or 50 nM), geldanamycin (100 or 200 nM), or a combination. **D)** Violin plots show the quantitation of colonies for LNCaP, LAPC4, and PC3 biological replicates (n=3 replicates in each experiment with 2 biological repeats). The asterisk indicates p<0.05.

**Fig 4:**
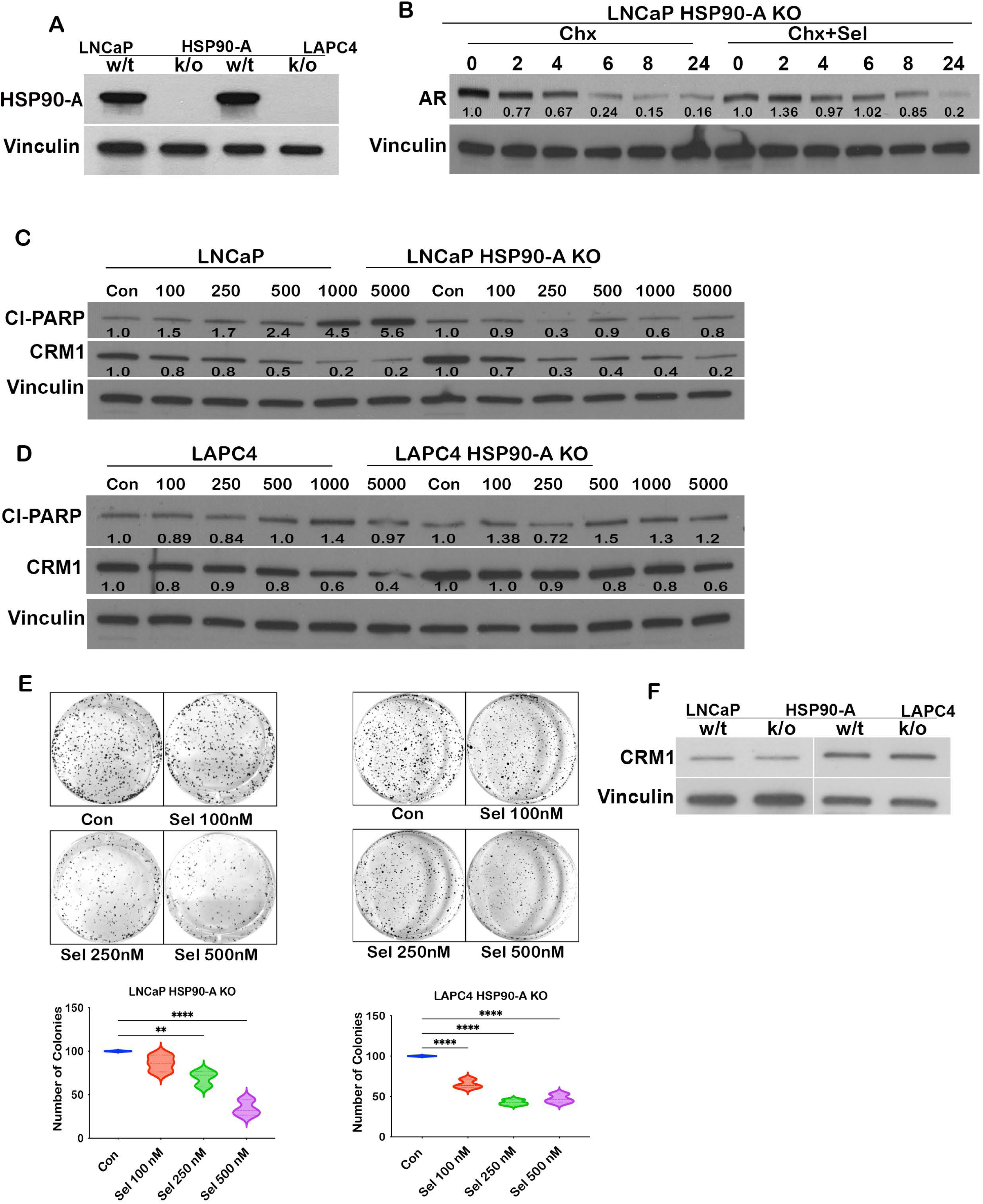
Effect of selinexor in HSP90α knockout cells. **A)** Western blot confirmation of HSP90α (90 kDa) knockout in LNCaP and LAPC4 cells. Vinculin was used as a housekeeper protein (124 kDa). B) AR protein stability in LNCaP HSP90α knockout cells post cycloheximide alone (100µM) or in combination with selinexor (1µM). Western blot showing cleaved PARP (89 kDa) and CRM1 (123 kDa) protein in LNCaP and LNCaP-HSP90α knockout cells **(C)** and LAPC4/LAPC4-HSP90α knockout cells **(D)** treated with dosage gradient of selinexor (0-5000 nM). **E)** Clonogenic survival assay for HSP90α knockout cells in LNCaP and LAPC4 treated with selinexor (0-500 nM) and quantitation of number of colonies shown as respective violin plots. **F)** Immunoblot to show relative levels of CRM1 protein (123 kDa) in wild-type LNCaP/ LAPC4 or HSP90α knockout cells. The asterisk on the violin plot indicates p<0.05.

### CRM1 inhibition affects DNA repair transcript and protein levels

In a previous study, we reported that doxorubicin, a topoisomerase II inhibitor, can potentiate selinexor-induced cytotoxicity [6]. Since AR regulates the transcription of several DNA repair genes (9), we next investigated whether CRM1 inhibition affects DNA repair gene expression and whether this is via AR. AR-positive (LNCaP, LAPC4, and VCaP) and AR-negative (PC3 and DU-145) prostate cancer lines were treated with varying concentrations of selinexor. RNA extracted after treatment was subjected to quantitative RT-PCR to evaluate transcript levels for various DNA repair genes of the homologous recombination (HR) and non-homologous end joining (NHEJ) DNA repair pathway **(Fig. 5A and Supp. Fig 1A)**. While LNCaP and VCaP AR-positive lines showed a decrease in the transcript level of most of the HR and NHEJ pathway genes tested, LAPC4 cells exhibited a dose-dependent upregulation of several HR and NHEJ pathway genes **(Fig. 5A)**. AR-negative PC3 cells did not exhibit much change in HR or NHEJ DNA repair genes upon CRM1 inhibition, while AR-negative DU-145 cells showed downregulation of several HR and NHEJ DNA repair genes at a higher dose **(Fig. 5A and Supp. Fig 1A)**. This data suggests that transcript levels of DNA repair pathway genes in response to CRM1 inhibition vary among cell types and may not depend on AR status. We next investigated whether inhibition of CRM1 affects DNA repair protein levels. Protein lysates from selinexor treated cells revealed that several proteins were differentially regulated upon selinexor treatment. Key proteins of the HR repair pathway (Rad51) and NHEJ repair pathway (Ligase IV) were consistently down upon CRM1 inhibition, irrespective of the AR status **(Fig. 5B and C and Supp. Fig 1B).** To ascertain whether the differential expression of genes upon CRM1 inhibition is irrespective of AR, we utilized a CRISPR-Cas9-engineered LNCaP-95 AR/AR v7 knockout cell line. Although AR null cells had lower levels of RAD51 and BRCA1 proteins than the parental cell type, as seen in Supp. Fig 1B, selinexor treatment decreased RAD51 protein and increased p53 expression in AR null cells. P53 protein increased in all cell types with p53 expression irrespective of whether the cells expressed wildtype or mutant (DU-145 cells) p53 **(Fig 5B and Supp Fig 1B).** This data suggests that there is a discordance between transcript and protein levels of DNA repair genes treated with selinexor. While the transcript level of RAD51 was increased in LAPC4 cells, RAD51 protein was decreased in all prostate cancer lines, including LAPC4 cells **(Fig. 5B and Fig S1B)**. Some of the repair proteins exhibited cell-type-specific responses to selinexor; this included NHEJ repair proteins like XLF, which was downregulated in PC3 and DU145 cells, and Ku-70, which was specifically downregulated in PC3 cells **(Fig. 5B and Fig S1B)**. Overall, this data suggested that both HR and NHEJ DNA repair pathways are affected by CRM1 inhibition. Intrigued by a dramatic decrease in RAD51 levels across all cell types, we investigated whether CRM1 plays a role in regulating RAD51 stability. LNCaP cells were treated with protein translation inhibitor cycloheximide alone or a combination of selinexor and cycloheximide, and western blots measured the decay of Rad51. As seen in **Figure 5D**, cells treated with a combination of cycloheximide and selinexor decreased the stability of the RAD51 protein. In order to gain insight into the mechanism of how CRM1 affects RAD51 protein stability, we treated cells with proteasome inhibitor MG132 or lysosomal inhibitor hydroxychloroquine or a combination of selinexor with the proteasomal or lysosomal inhibitor. As seen in **Figure 5E**, inhibition of the proteasomal pathway rescues the downregulation of RAD51 by selinexor, indicating that selinexor affects RAD51 protein stability by targeting RAD51 for proteasomal degradation.

**Fig 5:**
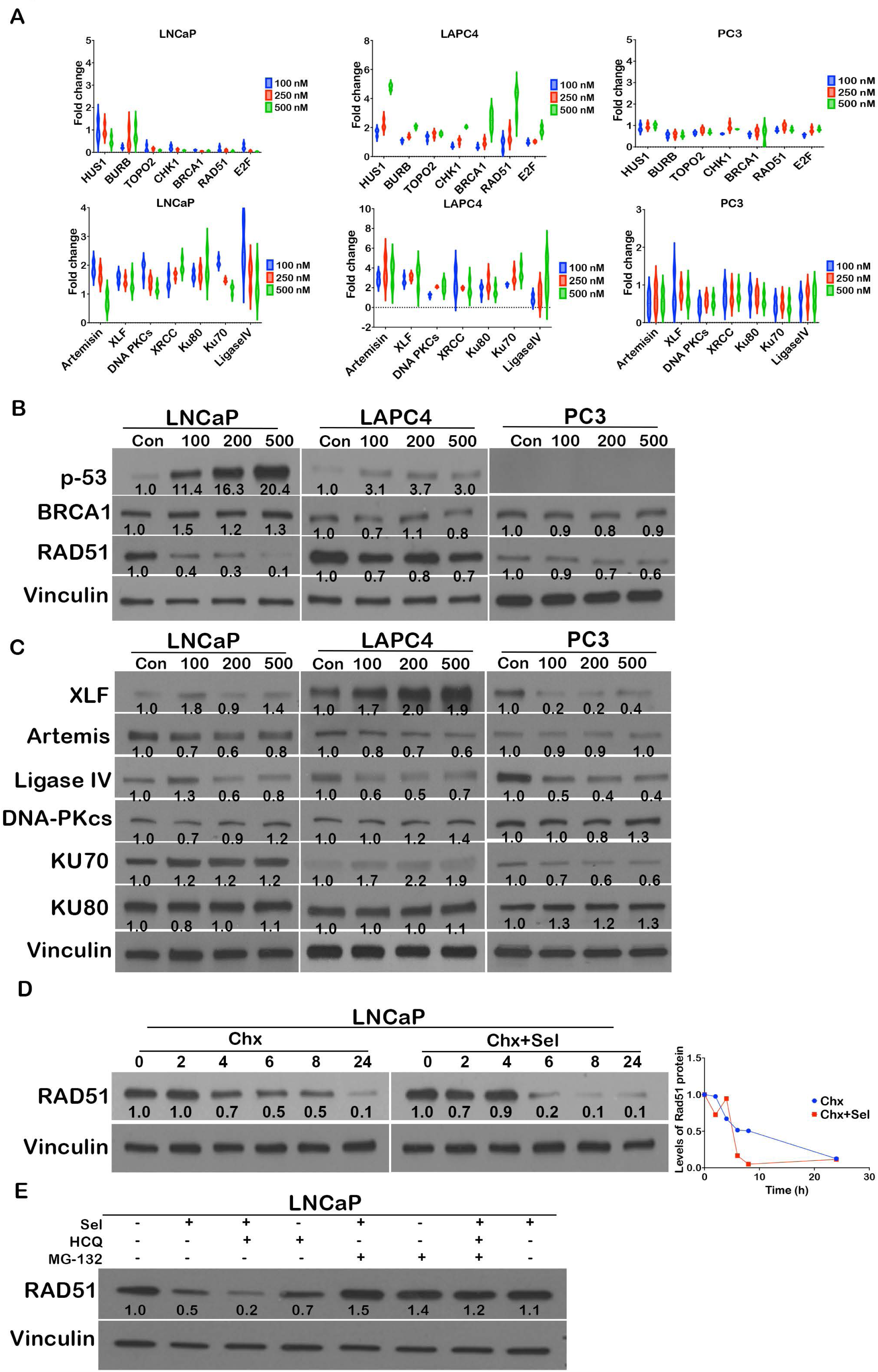
CRM1 inhibition affects DNA repair pathway proteins. **A)** Relative levels of transcripts for HUS1, BURB, TOPO2, CHK1, BRCA1, RAD51, and E2F for HR repair and Artemisin, XLF, DNA PKCs, XRCC, Ku80, Ku70, and LigaseIV for NHEJ DNA repair pathway in LNCaP, LAPC4, and PC3 cells. **B)** Protein levels of p53 (53kDa), BRCA1(220 kDa), RAD51(37 kDa) in LNCaP, LAPC4, and PC3 treated with selinexor (0-500 nM) for 48 hrs. **C)** Protein levels for XLF (39 kDa), Artemis (90 kDa), LigaseIV (100 kDa), DNA-PKcs (450 kDa), Ku70 (70kDa), and Ku80 (86 kDa) in LNCaP, LAPC4 and PC3 cells treated with selinexor (0-500 nM) for 48 hrs. **D)** Immunoblot showing the stability of RAD51 (37kDa) in LNCaP cells treated with cycloheximide (100µM) alone or in combination with selinexor (1µM). Line graph showing normalized levels of RAD51 protein. **E)** Western blot analysis of RAD51 protein in LNCaP cells treated with selinexor (1µM), MG132 (10 µM), or hydroxychloroquine (50 µM). Vinculin (124kDa) was used as a housekeeper protein for normalization (B-E).

### CRM1 affects DNA repair function in prostate cancer cells

Our data suggests that CRM1 affects the stability of key DNA repair enzymes. To evaluate the functional consequence of CRM1 inhibition on the kinetics of DNA repair, we first studied whether CRM1 inhibition has an effect on the activation of ATM, a central DNA repair kinase, and p53, the central guardian of the genome. LNCaP cells were irradiated with 4Gy of radiation in the presence and absence of CRM1 inhibition. Cell lysates were collected at different time points, and activation of ATM or p53 was evaluated using antibodies against activated forms of respective proteins. While there was a slight decrease in activated ATM upon CRM1 inhibition, activation of p53 was markedly delayed upon CRM1 inhibition **(Fig. 6A and Supp. Fig 2A)**. This suggested that inhibition of CRM1 delays the kinetics of repair protein activation. We next investigated whether inhibition of CRM1 might alter the recruitment of BRCA1 or RAD51 proteins through immunofluorescence. We did not find any change in the recruitment of these proteins upon CRM1 inhibition **(Supp. Fig 2B)**. Our analysis showed that CRM1 has several S/TQ phosphorylation motifs that could potentially be phosphorylated by ATM/ATR kinases **(Fig. 6B)**. A number of these motifs are present as surface residues on CRM1. To investigate whether CRM1 can be phosphorylated upon DNA damage, we utilized a phospho-Tag gel on lysates that have been irradiated. Phospho-tag gels can discern the phosphorylated form of the protein as mobility shifts (10). While CRM1 had various protein bands, indicating phosphorylation at basal state, upon irradiation, there was increased phosphorylation of several residues **(Fig. 6C)**. This suggests that CRM1 may function as a DNA repair effector, facilitating repair of damaged DNA. One of the hallmarks of many DNA repair proteins is their ability to localize to DNA double-strand breaks. To investigate whether CRM1 can localize to DNA double-strand breaks, we irradiated LNCaP cells with 4Gy radiation and co-stained for CRM1 and γ-H2AX, a surrogate marker of DNA double-strand breaks. As seen in **Fig. 6D**, while CRM1 is localized to the nuclear envelope and the nucleoplasm in non-irradiated cells, in irradiated cells, CRM1 is partially localized to DNA double-strand breaks. We confirmed this using etoposide, which induces double-strand DNA breaks, and p53bp1, another surrogate marker of DNA double-strand breaks. We observed CRM1 strongly colocalized with p53BP1 on chromatin strands and partially to DNA repair foci **(Fig. 6E)**. This suggests that CRM1 may have a more direct role in DNA repair as an effector protein. To investigate whether inhibition of CRM1 affects the DNA repair capacity of prostate cancer cells, we utilized an γ-H2AX clearance assay to quantify DNA repair in irradiated cells treated with selinexor. While control cells had fewer γ-H2AX foci after 4h of repair, selinexor-treated cells had an increased number of unrepaired foci **(Fig. 6F)**. This suggested that inhibition of CRM1 results in a decrease in DNA repair in prostate cancer cells. This was confirmed with an alkaline comet assay that quantifies DNA double-strand breaks as tail lengths **(Fig. 6G)**. We also noted an increase in the number of micronuclei in irradiated cells that we treated with selinexor **(Fig. 6H).** Micronuclei result from chromosomal instability and have been linked to faulty homologous recombination DNA repair (11). To investigate whether inhibition of CRM1 affects HR repair, we performed a fluorescent protein-based homologous recombination assay (12). As seen in **Figure 7A and Supp. Fig 2C**, inhibition of CRM1 decreased HR repair in LNCaP cells dose-dependently. To find whether inhibition of CRM1 would potentiate any of the preclinical or clinical DNA repair pathway inhibitors, we treated LNCaP cells with selinexor alone and in combination with various DNA repair pathway inhibitors (ATR inhibitor VE821, ATM inhibitor AZ32, PARP inhibitor Olaparib, and DNA-PK inhibitor M3814). As seen in **Figures 7B and C and Supp. Fig 2D and E**, selinexor enhanced growth inhibition of AZ32 in LNCaP cells and Olaparib in DUI145 cells. Combination with other DNA repair pathway inhibitors showed no significant growth reduction and, intriguingly, showed a tendency towards cytoprotection. This data suggests that CRM1 inhibitions may be combined with only certain DNA repair pathway inhibitors, and the effect is cell-type specific.

**Fig 6:**
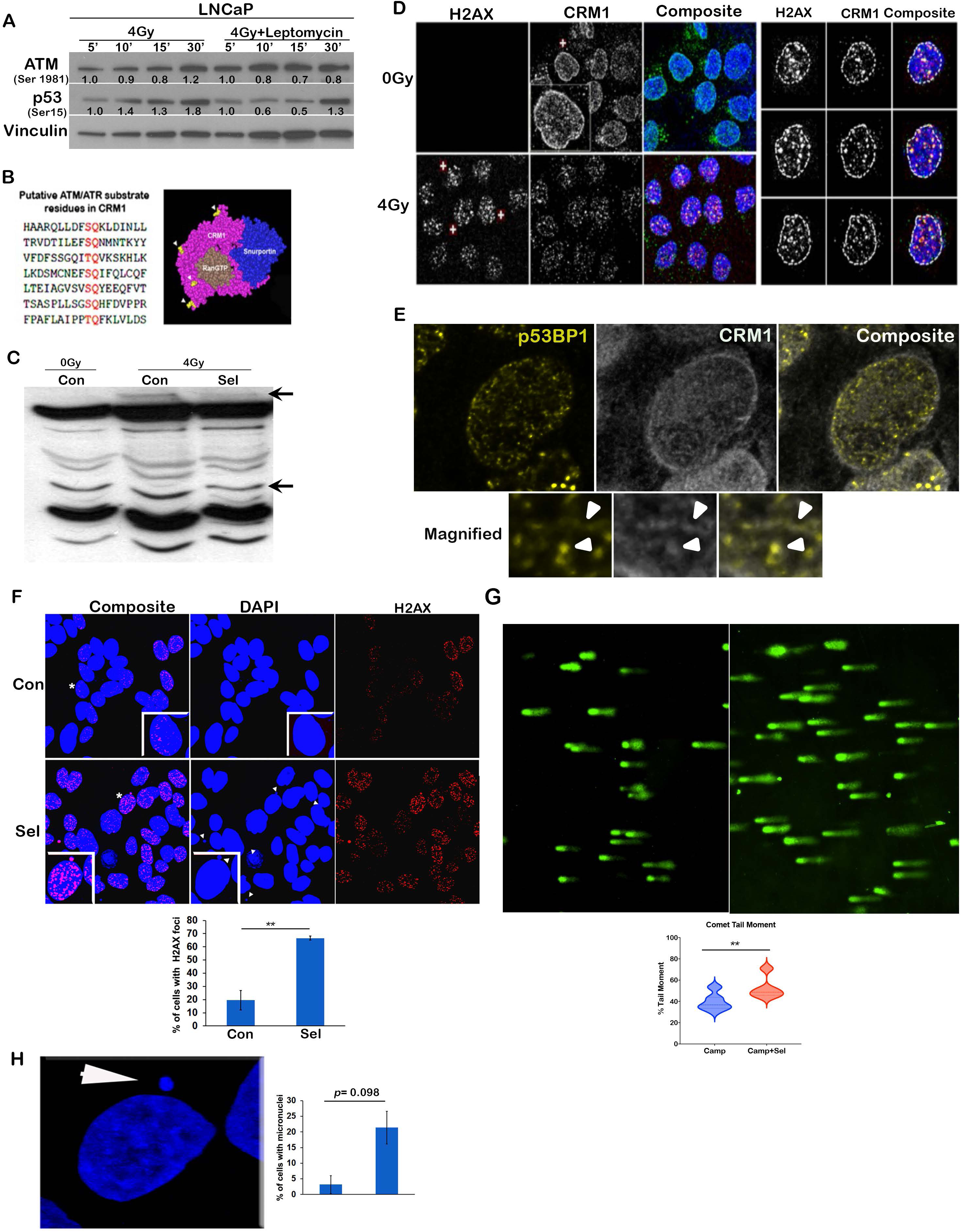
CRM1 inhibition affects DNA repair in prostate cancer cells. A) Western blot showing phosphorylated levels of ATM kinase (350 kDa) and p53 protein (53 kDa) in 0 or 4 Gy radiated LNCaP cells treated with either vehicle or Leptomycin B (10nM) for the indicated time. Vinculin was used as housekeeper control (124 kDa). B) Various S/T-Q residues (serine/threonine followed by glutamine) in CRM1 protein and their localization on CRM1 protein to indicate putative phosphorylation sites. C) Phosphotag analysis of CRM1-Flg protein expressed using a plasmid construct in LNCaP cells. Western blot showing various phosphorylated forms of CRM1 probed using an anti-Flg antibody. Samples were resolved using a modified polyacrylamide gel to retard the mobility of phosphorylated protein moieties. Arrows indicate differentially appearing intensities of some of the CRM1 phospho-forms. D) Representative confocal microscopy images of localization of damaged DNA makers H2AX and CRM1 and colocalization analysis. E) Localization of p53 BP1 with CRM1 in LNCaP cells treated with etoposide for 12h followed by 1h of repair. Lower images are magnified views of a region from the above panel. Arrowheads depicts CRM1 and p53BP1 colocalization. F) Levels of damaged DNA in vehicles or selinexor-treated LNCaP cells using confocal microscopy and its quantitation using image analysis on ImageJ software. G) Comet assay analysis for LNCaP cells treated with either vehicle or selinexor. The tail moment was calculated using ImageJ and plotted on a violin plot. H) DAPI staining to estimate the micronuclei in LNCaP cells treated with selinexor.

**Fig 7:**
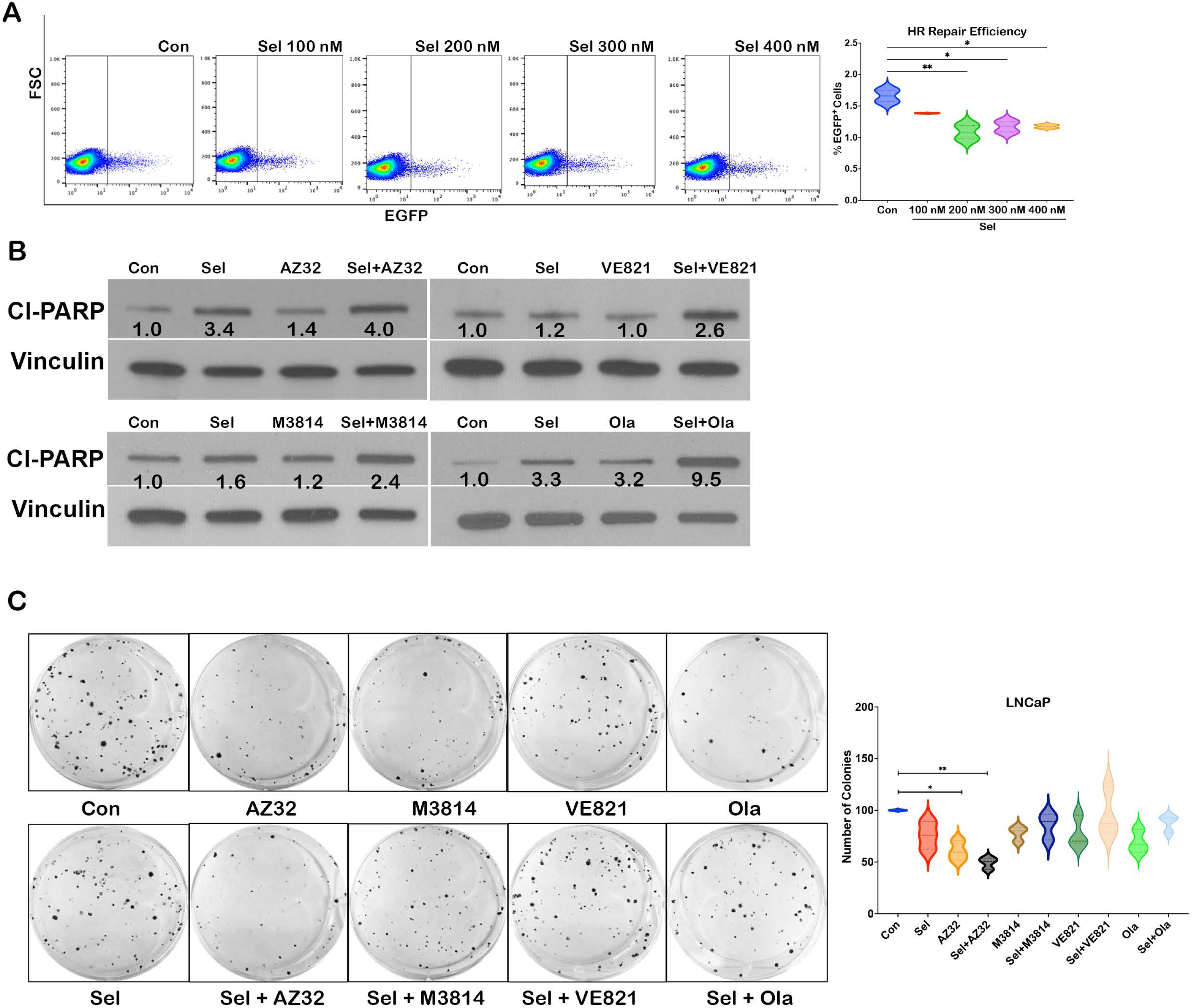
DNA repair pathways can be targeted with CRM1 inhibitions. A) Flowcytometry-based analysis of LNCaP cells treated with selinexor for homologous recombination-mediated DNA repair using a mutant EGFP construct (pJustin) and recovery of green fluorescence indicated HR repair. The violin plot shows the quantitation of the EGFP fluorescence and compares LNCaP cells treated with different dosages of selinexor (100-400 nM) for 12 hrs. The asterisks indicate p<0.05. B) Western blot analysis of LNCaP cells treated with either vehicle or single agent selinexor (100 nM) or AZ32 (3.1 µM) VE821 (6.12 µM) M3815 (0.2 µM) or Olaparib (5 µM) or indicated combinations for 48 hrs. Protein samples were probed with cleaved PARP (89 kDa) or housekeeper protein vinculin (124 kDa). C) Clonogenic assay to test long-term fitness of LNCaP cells treated with selinexor, AZ32, VE821, M3814, or Olaparib as a single agent or combination. Violin plot showing quantitation of the number of colonies in each treatment type. The asterisks indicate p<0.05.

## Discussion

Our data suggests that CRM1 affects AR mRNA and protein stability; this dual control over AR mRNA and protein results in the loss of AR protein upon CRM1 inhibition, affecting its function. HuR is known to interact with and positively affect the stability of the AR mRNA [25], and cytoplasmic HuR correlates with prostate cancer aggressiveness [26, 27]. Our data suggests that inhibition of CRM1 prevents its nuclear-cytoplasmic shutting, restricts HuR to the nucleus, and decreases AR mRNA stability. We further present evidence suggesting that AR protein stability may be affected by CRM1, independent of its effect on AR mRNA stability. AR protein is known to be stabilized by AR chaperones that keep AR in a poised ligand-binding state. Targeting AR chaperones clinically with small molecules to inhibit prostate cancer growth has met with mixed results primarily due to toxicities associated with chaperone inhibitors [28–30]. Our data suggests that HSP90 alpha harbors a nuclear export signal and is exported to the cytoplasm in a CRM1-dependent manner. Inhibiting CRM1 affects HSP90 nuclear export, thereby decreasing cytoplasmic HSP90 that is needed for AR protein stability. Combining CRM1 inhibition with HSP90 inhibitor potently reduces prostate cancer cell growth and survival. Intriguingly, this combination is not selective for AR-positive prostate cancer cells but is equally potent against AR-negative prostate cancer, likely as CRM1 may affect the export of other oncogenic driver proteins that are HSP90 client proteins besides AR. Future preclinical work in mouse models and patient-derived xenograft models would provide more insights into whether this combination would be effective against prostate cancer growth, irrespective of AR status. Our data shows that cells that have decreased HSP90 alpha may resist CRM1 inhibitors. Although the mechanism for this resistance couldn’t be elucidated from our data, it is likely to involve apoptosis resistance mechanisms given the role of HSP90 alpha in regulating DNAPK activation and apoptosis [31]. Since AR is known to be involved with the transcription of genes of DNA damage and response pathway [11, 12], we hypothesized that downregulation of AR function upon AR inhibition might affect the expression of DNA damage response pathway genes, making it vulnerable to DNA repair pathway inhibitors. We found that CRM1 affects RNA and protein levels of several DNA repair proteins that function in the NHEJ or HR pathway, irrespective of the AR status. Rad51, a key HR protein, was downregulated upon CRM1 inhibition across all prostate cancers. This suggested that DNA repair, specifically HR repair, may be a key vulnerability upon CRM1 inhibition. Inhibition of CRM1 combined with ATR, ATM, DNA PK, or PARP inhibitor increased apoptosis in the short term. However, long-term clonogenic survival assays for all inhibitors except AZ32 (ATM inhibitor) resulted in cytoprotection compared to single-agent treatment. We were surprised to find the combination treatment of CRM1 inhibitor with PARP inhibitor Olaparib, known to decrease survival in the presence of HR repair deficiency, ineffective in reducing clonogenic survival in all cell types except DU145 cells. Our results contrast other reports that combined CRM1 inhibitors with Olaparib and found an enhanced decrease in the clonogenic survival [32]. One of the reasons could be the difference in dose and culture conditions. However, it is also likely that a combination treatment of CRM1 inhibitors with inhibitors of certain DNA repair pathways may trigger pathways that benefit long-term clonogenic survival. In summary, our data suggests that CRM1 can affect prostate cancer growth by regulating stability of androgen receptor and impacts DNA repair in prostate cancer cells independent of the androgen receptor. A combination of DNA repair pathway inhibitors along with CRM1 inhibitors may be an effective strategy in decreasing prostate cancer growth; however, the choice of repair pathway that is inhibited may dictate the therapeutic outcome.

## Acknowledgements

The work was supported by W81XWH1910724, 1R01CA243184, Patrick C Walsh, and PCF Challenge awards. DoD grants W81XWH2210118, HT94252310029, and PCF Young Investigator Award 21YOUN22 support RK. We also acknowledge Dr. Paschal Bryce for providing the plasmid construct for AR-GFP expression.

---

**Supplementary figure 1:** DNA repair pathway proteins in VCaP and DU145 cells. A) Relative levels of transcripts for HUS1, BURB, TOPO2, CHK1, BRCA1, RAD51, and E2F for HR repair and Artemisin, XLF, DNA PKCs, XRCC, Ku80, Ku70, and LigaseIV for NHEJ DNA repair pathway in VCaP and DU145 cells. B) Protein levels of p53 (53kDa), BRCA1(220 kDa), RAD51(37 kDa, XLF (39 kDa), Artemis (90 kDa), LigaseIV (100 kDa), DNA-PKcs (450 kDa), Ku70 (70kDa), and Ku80 (86 kDa) in in VCaP, and DU145 treated with selinexor (0-500 nM) for 48 hrs.

**Supplementary Figure 2:** A) Western blot showing phosphorylated p53 (Serine 15) (53 kDa) in a vehicle or selinexor-treated LNCaP radiated with 4Gy radiation at indicated time points. Vinculin (124 kDa) was used as a housekeeper protein for normalization. B) Representative photomicrograph from confocal microscopy showing localization of Rad51 and BRCA1 on damaged DNA probed with γH2AX. DAPI was used to stain the nuclei. C) Inverted microscope captured images of LNCaP cells expressing pJustin plasmid (a mutant form of EGFP, which produces fluorescence upon DNA repair), treated with either vehicle or indicated doses of selinexor. Bar graphs indicate the quantitation of the number of cells showing green fluorescence. The asterisks indicate p<0.05. D) Immunoblotting of proteins isolated from AR negative PCa, DU145, treated with vehicle or single agent selinexor, AZ32, VE821, M3814, or Olaparib, or indicated combinations. Cleaved PARP (89 kDa) was used as an apoptosis indicator, and vinculin (124 kDa) was used as a housekeeper protein for normalization. E) Clonogenic assay for DU145 treated with conditions mentioned in D), representative images for stained colonies are shown, and violin plot shows quantitation of the average number of colonies from replicate dishes. The asterisks indicate p<0.05.

## Notes

### Competing Interest Statement

The authors have declared no competing interest.

