## Supplemental Fig 1-2 and Table 1 for "CRM1 regulates androgen receptor stability and impacts DNA repair pathways in prostate cancer, independent of the androgen receptor"

### Supplimentary Figure 1

A

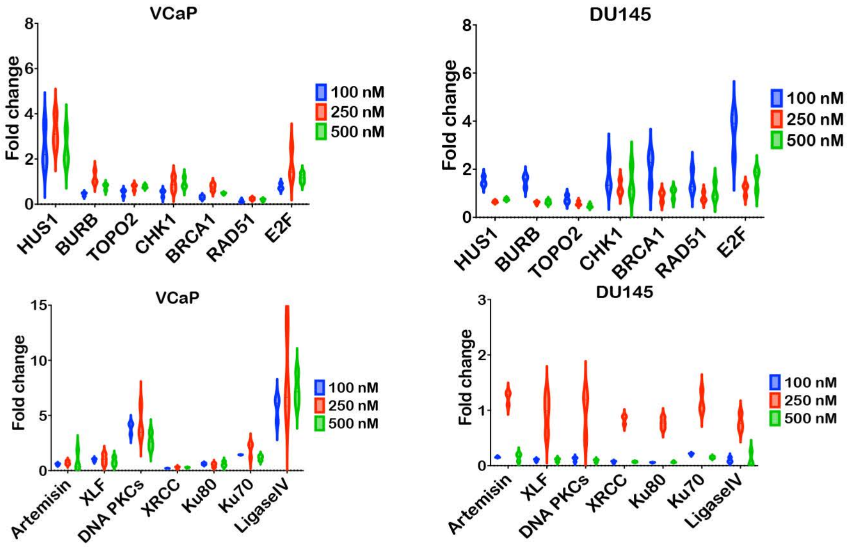


B

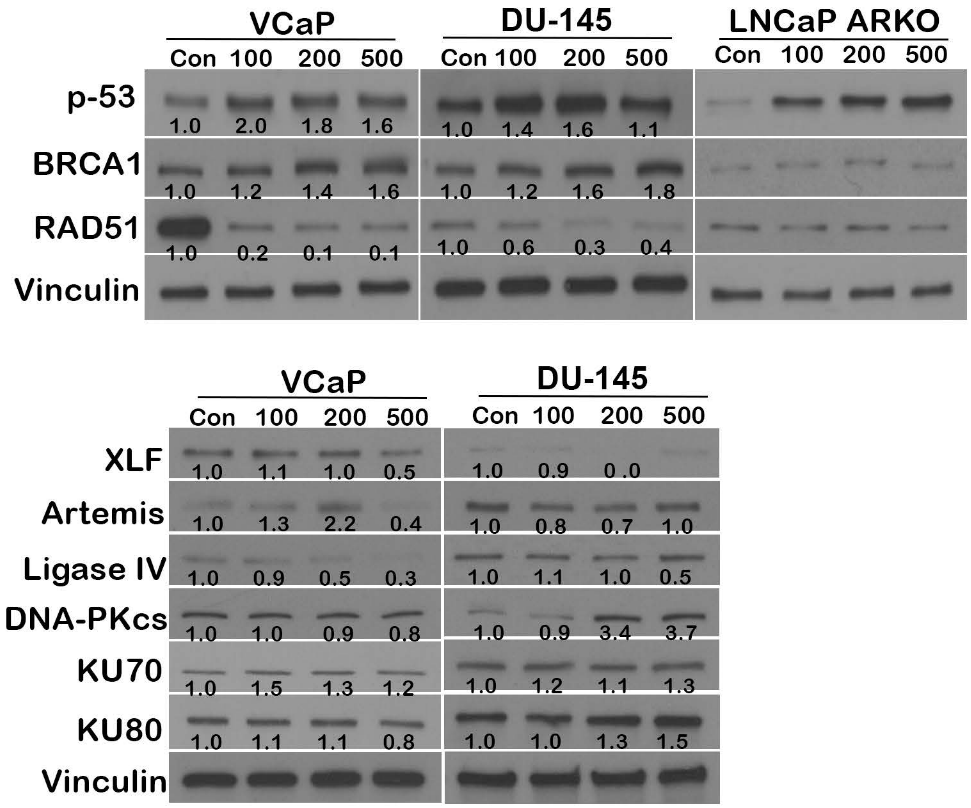

#### Supplimentary Figure 2

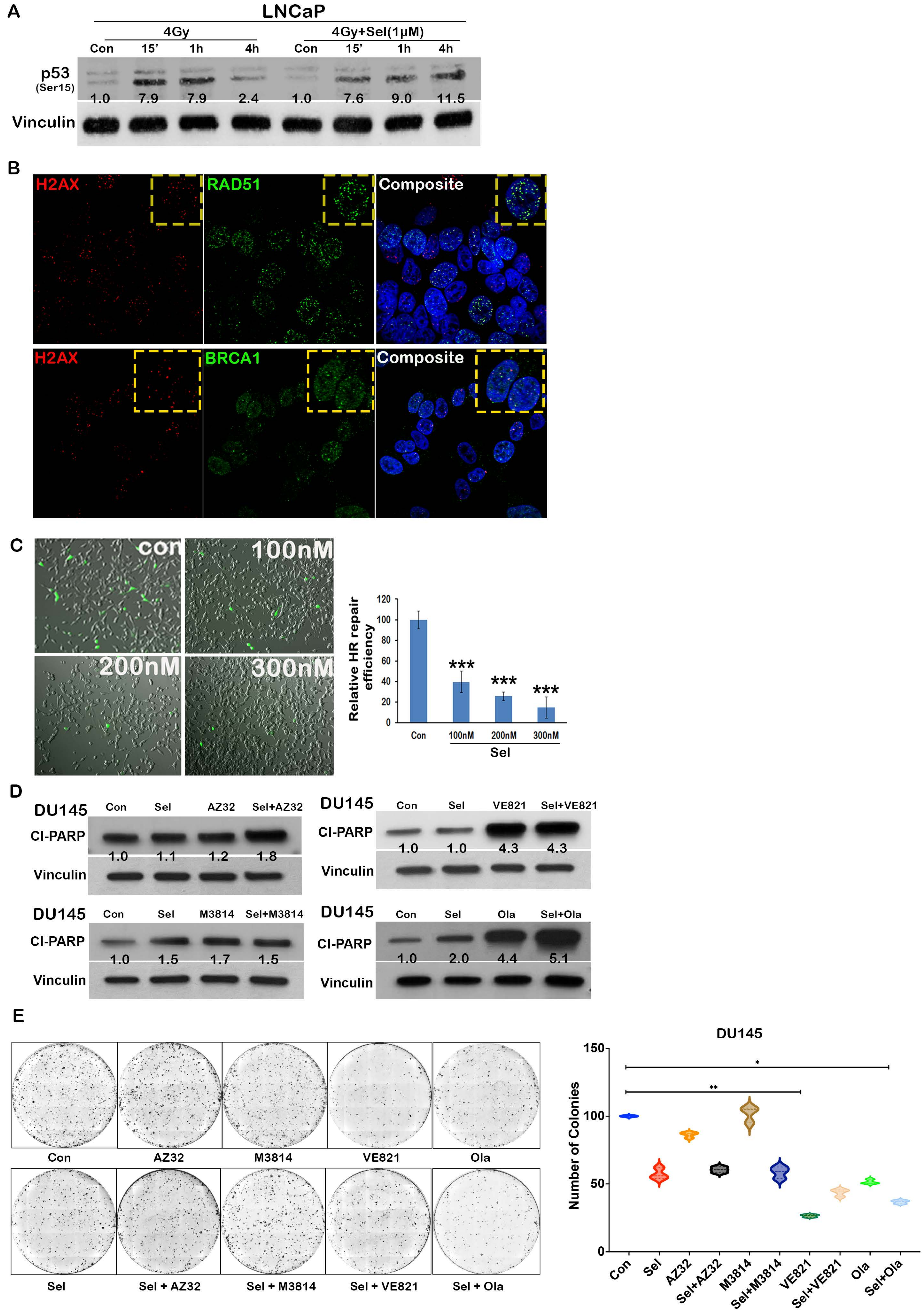

**Supplementary table 1**: List of Antibodies

| S No | Antibody | Cat No | Species | Manufacturer |
|---|---|---|---|---|
| 1 | AR | sc-7305 | Mouse | Santa Cruz Biotechnology |
| 2 | AR v7 | sc-272739 | Mouse | Santa Cruz Biotechnology |
| 3 | Actin | A5316 | Mouse | Sigma |
| 4 | Cleaved PARP | 9541S | Rabbit | Cell Signaling Technology |
| 5 | Vinculin | 05-386 | Mouse | Millipore Sigma |
| 6 | HSP90 A | ADI-SPA.8400 | Rat | Enzo |
| 7 | CRM1 | 47249S | Rabbit | Cell Signaling Technology |
| 8 | p53 | OP43-100UG | Mouse | CalBiochem |
| 9 | BRCA1 | sc6954 | Mouse | Santa Cruz Biotechnology |
| 11 | RAD51 | 88755 | Rabbit | Cell Signaling Technology |
| 11 | XLF | 2854 | Rabbit | Cell Signaling Technology |
| 12 | Artemis | 13381 | Rabbit | Cell Signaling Technology |
| 13 | Ligase IV | 14649 | Rabbit | Cell Signaling Technology |
| 14 | DNA PKcs | 38168 | Rabbit | Cell Signaling Technology |
| 15 | KU70 | 4588 | Rabbit | Cell Signaling Technology |
| 16 | KU80 | 2180 | Rabbit | Cell Signaling Technology |
| 17 | ATM SER 1981 | 50-199-0529 | Mouse | Rockland Immunochemicals |
| 18 | ATM | 2873S | Rabbit | Cell Signaling Technology |
| 19 | p53 Ser 15 | 9286S | Mouse | Cell Signaling Technology |
| 20 | Flg | F3165 | Rabbit | Sigma |
| 21 | HuR | Sc5261 | Mouse | Santa Cruz Biotechnology |
| 22 | p53BP1 | | | |
| 23 | $\gamma$-H2AX | ZMS05636 | Mouse | Millipore Sigma |
| 24 | Anti-Mouse HRP | 7076S | Horse | Cell Signaling Technology |
| 25 | Anti-Rat HRP | 7077S | Goat | Cell Signaling Technology |
| 26 | Anti-Rabbit HRP | 7074S | | Cell Signaling Technology |

| | | | | |
|---|---|---|---|---|
| 27 | Goat anti-Mouse IgG (H+L) Highly Cross-Adsorbed Secondary Antibody, Alexa Fluor 555 | A21424 | Goat | Thermo Fisher Scientific |
| 28 | Goat anti-Mouse IgG (H+L) Highly Cross-Adsorbed Secondary Antibody, Alexa Fluor 488 | A11029 | Goat | Thermo Fisher Scientific |
| 29 | Goat anti-Rabbit IgG (H+L) Highly Cross-Adsorbed Secondary Antibody, Alexa Fluor 488 | A11034 | Goat | Thermo Fisher Scientific |
| 30 | Goat anti-Rabbit IgG (H+L) Highly Cross-Adsorbed Secondary Antibody, Alexa Fluor 555 | A21429 | Goat | Thermo Fisher Scientific |
